# Artificial endoplasmic reticulum-lipid droplet tethers facilitate lipid incorporation into lipid droplets

**DOI:** 10.64898/2026.06.25.734520

**Authors:** Victoria H. Williams, Gregory E. Miner, Sarah Cohen

## Abstract

Lipid droplets (LDs) are ubiquitous organelles that store neutral lipids to meet cellular energetic and signaling needs. As a unique monolayer structure, LDs arise from the endoplasmic reticulum (ER) and acquire proteins and lipids through their membrane contact sites (MCSs) with the ER. In this study, we exogenously induce ER-LD MCSs using a dimerization-dependent fluorescent protein (ddFP) system. Strikingly, inducing these MCSs increases LD size without influencing LD total amount per cell, in a manner that is distinct from LD biogenesis induced by the dietary fatty acid oleic acid. By examining the trafficking of the triacylglycerol synthesis enzyme DGAT2 under ddFP induction, we found that artificial tethering recruits LD proteins to the ER-LD interface but not to the LD surface, unlike oleic acid supplementation. However, by supplementing ddFP-transfected cells with fluorescent fatty acids, we found that ddFP-positive LDs preferentially incorporate exogenous lipid, suggesting that inducing MCSs can facilitate ER-to-LD lipid transfer. These results demonstrate ddFPs as a tool for manipulating LD MCSs and elucidate the role of ER-LD MCSs following LD biogenesis to continue to promote LD growth.

## Introduction

Lipid droplets (LDs) are ubiquitous structures seen in both prokaryotes and eukaryotes (Farese and Walther, 2025; Waltermann et al, 2005). The LD consists of a neutral lipid core surrounded by a phospholipid monolayer, a membrane structure that is unique among cytosolic membrane-bound organelles (Farese and Walther, 2025). While initially considered inert lipid storage units, LDs within eukaryotes have gained additional attention as *bona fide* organelles with a distinct proteome, that undergo biogenesis, fusion, and turnover (reviewed in: Olzmann and Carvalho, 2019; Henne and Cohen, 2026). LDs can sequester potentially lipotoxic lipid species within neutral lipids and liberate stored fatty acids for beta oxidation by the mitochondria, involving LD-mitochondria membrane contact sites (MCSs) for both processes depending on cell type and condition (Rambold et al, 2015; Benador et al, 2019, Miner et al, 2023). LDs also serve as the source for key signaling lipids and have emerging roles in both viral replication and cellular immunity (reviewed in Kaiser et al, 2022; Zadoorian et al, 2023). The endoplasmic reticulum (ER) is the major site of intracellular lipid synthesis; LDs begin as accumulations of neutral lipids, such as triacylglycerols (TAGs), between the leaflets of the ER (reviewed Walter et al, 2017). Key LD biogenesis proteins, such as seipin, help mediate the process of forming LDs from this oil lens, regulating LD size and protein recruitment (Cartwright et al, 2014). Seipin oligomerizes at the ER-LD interface and mediates the partitioning of neutral lipid from the ER to nascent LDs, allowing for the expansion of smaller LDs (Arlt et al, 2022; Wang et al, 2016). In contrast, without seipin, larger LDs tend to increase in size at the expense of smaller LDs in a process known as ripening (Szymanski et al, 2007; Fei et al, 2008; Salo et al, 2019).

Proteins traffic to the LD either through recruitment from the cytoplasm (cytoplasm-to-LD, CYTOLD pathway) or by moving from ER membranes to the LD (ER-to-LD, ERTOLD pathway) (Olarte et al, 2022). Because LDs are surrounded by a phospholipid monolayer, neither of these targeting methods happen through the well-characterized vesicular membrane trafficking pathways of membrane-bound organelles, although proteins involved in these processes have been shown to play roles in LD targeting, suggesting an alternative role for such proteins in the context of trafficking to monolayer organelles (Walther and Farese, 2025). Previous studies suggest that ERTOLD targeting occurs through two different types of ER-LD contact sites – those early in LD biogenesis largely regulated by seipin and its interactors, and a less well-characterized type of ER-LD contact that allows for later targeting of additional ERTOLD proteins, largely studied in *Drosophila* cells (Song et al, 2022).

MCSs are difficult to resolve by diffraction limited light microscopy, because they are defined as sites of close apposition between two organelles (often 10-30 nm between membranes), mediated by tether proteins (reviewed in Calì et al, 2025; Prinz et al, 2020; Scorrano et al, 2019). Dimerization-dependent fluorescent proteins (ddFPs) utilize the intrinsic dimerization of red fluorescent proteins to effectively label two proteins of interest when they are in proximity in live cells, allowing for the definitive identification of MCSs. ddFPs generate increased fluorescence at MCSs when a weakly fluorescent “A” monomer interacts with a nonfluorescent “B” monomer (Alford et al, 2012a, Alford et al 2012b, Ding et al, 2015). This approach has been used to study a variety of intracellular organelle interactions such as ER-mitochondria and other contacts (Abrisch et al 2020; Alford et al, 2012b; Lee et al, 2020; Naon et al, 2016; Nguyen and Voeltz, 2022). ddFPs offer the advantage of allowing the user to visualize MCSs in live cells using widely available confocal microscopy rather than more specialized super-resolution techniques. Recently, a suite of tools to visualize MCSs at 7 distinct organelles using ddFPs that can be combined pairwise has been created, allowing for the study of a wide variety of MCSs in live cells (Miner et al, 2024).

While ddFPs have relatively weak affinity for one another, here we show that at higher expression levels the system can induce MCSs between the ER and LDs. We leveraged the induction of MCSs by this system as an artificial tether to manipulate ER-LD MCSs without modifying expression of known ER-LD MCS tether proteins. We sought to parse which attributes of ER-LD MCSs are specifically dependent on the activity or structural features of endogenous ER-LD MCS proteins and which were functions of retained access of mature LDs to ER pools of lipid and proteins which could be manipulated using artificial tethers. We found that inducing ER-LD contacts modulated distribution of cellular lipid into LDs and increased LD size, in a way that differed from fatty acid supplementation. We also found that ddFP-positive LDs were larger than ddFP negative LDs. We hypothesized that ddFP tethers increase LD size by sustaining ER-LD MCSs and performing known functions of ER-LD MCSs, either protein recruitment to the LD surface or direct movement of lipids synthesized in the ER to LD. We found that the TAG synthesis enzyme diacylglycerol O-acyltransferase 2 (DGAT2), which translocates from the ER to LDs upon fatty acid supplementation, does not fully translocate in ddFP-transfected cells but instead retains ER localization and traffics to the ER-LD interface of ddFP-positive LDs. In contrast, fluorescent fatty acid supplementation with FL-C12 revealed that the storage of cellular fatty acid is biased towards ddFP-positive LDs, increasing their size over time. Together, we demonstrate that ddFP tools are not only useful in visualizing MCSs, but can also be leveraged to exogenously manipulate MCSs in ways that shift cellular lipid storage. Our data also suggest that such artificial contacts can perform some, but not all, of the functions of known ER-LD tether proteins.

## Results

### Dimerization-dependent fluorescent proteins effectively localize to organelles of interest

While the original ddFP constructs were optimized for the green A (GA) fluorescent monomer at LD-organelle contact sites (Miner et al, 2024), we sought to further optimize the use of and confirm the localization of the red A (RA) monomer version to MCSs of interest. To label LDs, we used the LD-B monomer, which we previously tested and found effective in labeling a variety of MCSs when combined with GA monomers targeted to other organelles. The LD-B construct targets the non-fluorescent ddFP monomer to LDs using the hairpin domain of GPAT4. The ER-RA construct utilizes the transmembrane domain of CYP2C1 to target the weakly fluorescent RA monomer to the cytoplasmic face of the ER. We confirmed that both the ER-RA and LD-B monomers correctly co-localized with markers for their designated organelles of interest when transfected with corresponding cytoplasmic ddFP constructs, Cyto-B and Cyto-RA respectively (Figures 1A-C). We observed colocalization of the dimerized ER-RA/Cyto-B signal with the ER organelle marker that was distinct from the pattern of the LD dye BODIPY493 (Figure 1B). We also confirmed increased fluorescence intensity of dimerized LD-B/Cyto-RA around LDs that was distinct from the ER pattern (Figure 1C). We then transfected ER-RA and LD-B together and found that the dimerized monomers brightly label sites of close apposition between the two organelles of interest (Figure 1D). Crucially, we also observed specificity in our label; while some LDs displayed complete rings of ER surrounding the LD, others showed partial rings or no ddFP labelling, suggesting that our label can distinguish between LDs with and without ER-LD contacts and can identify specific regions of contact on an LD of interest.

**Figure 1:**
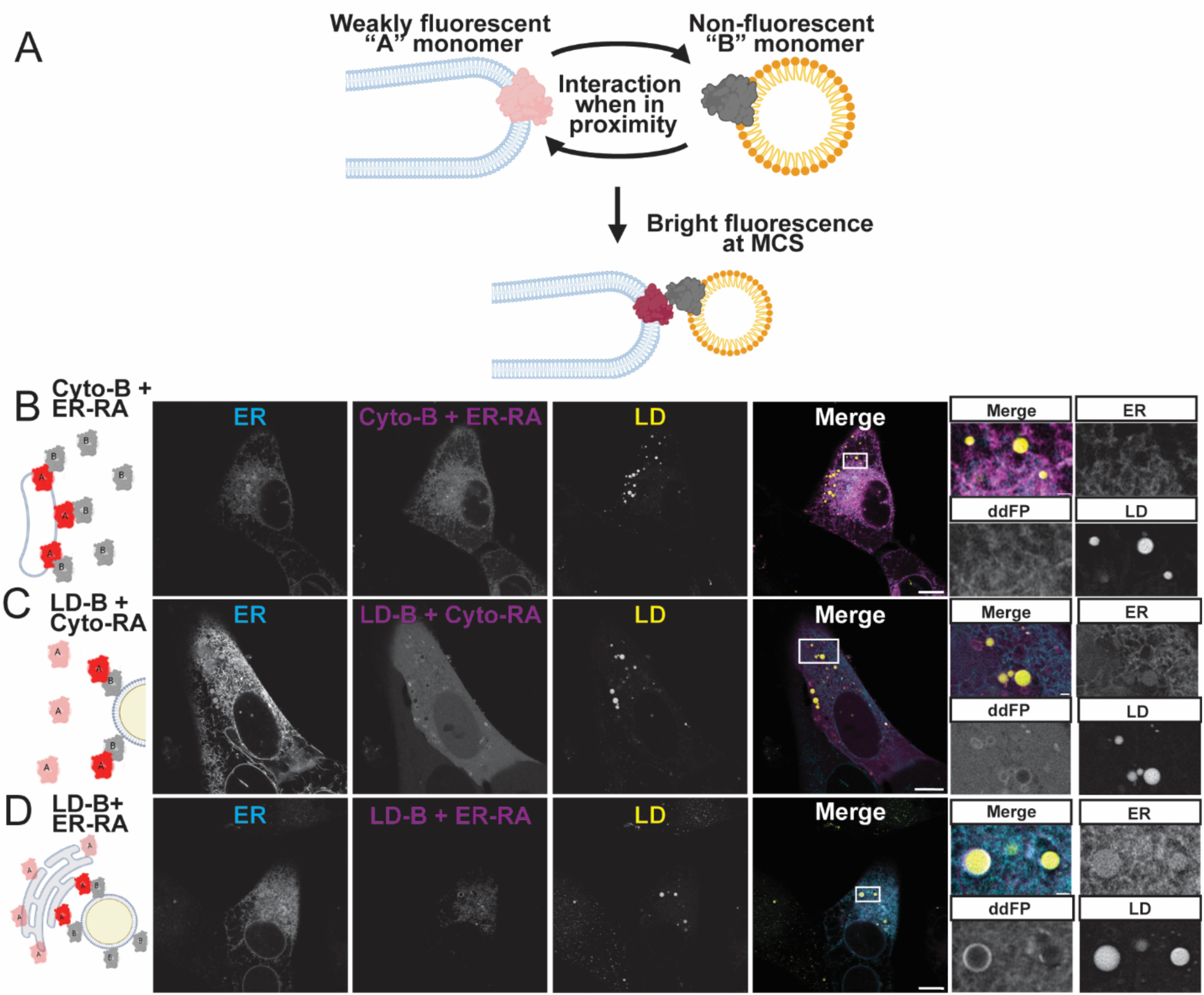
Contact-FP labels ER-LD contact sites. A) Schematic for Contact-FP dimerization-dependent fluorescent protein system. B) ER-RA correctly localizes to ER in U-2 OS. ER-RA plasmid and cytoplasmic-B plasmid were transfected with BFP-KDEL to label ER. BODIPY493 dye was added prior to imaging to label LD. C) LD-B correctly localizes to LD in U-2 OS. ER-RA plasmid and cytoplasmic-RA plasmid were transfected with BFP-KDEL and LD labelled with BODIPY493. D) Contact-FP labels ER-LD Contact sites. Scale bars: 10 µm.

### Inducing ER-LD MCSs with ddFPs increases lipid droplet size

We next confirmed whether we could induce interaction between the ER and LDs by expressing ddFPs. ddFPs have been previously titrated down to visualize LD-mitochondria dynamics with minimal perturbation or expressed at higher levels to induce LD-mitochondria contacts (Miner et al, 2024). Indeed, we found that at the amount of plasmid transfected in this study, overlap of the markers for ER and the LD dye increased compared to either ddFP monomer transfected separately, increasing the percent area of segmented LD object overlapping ER (Figures 2A-C).This suggests that our constructs do induce ER-LD MCSs at these concentrations and confirms that ddFPs can be used not only to visualize MCSs, but also to artificially manipulate MCSs by inducing exogenous tethering of organelles of interest.

**Figure 2:**
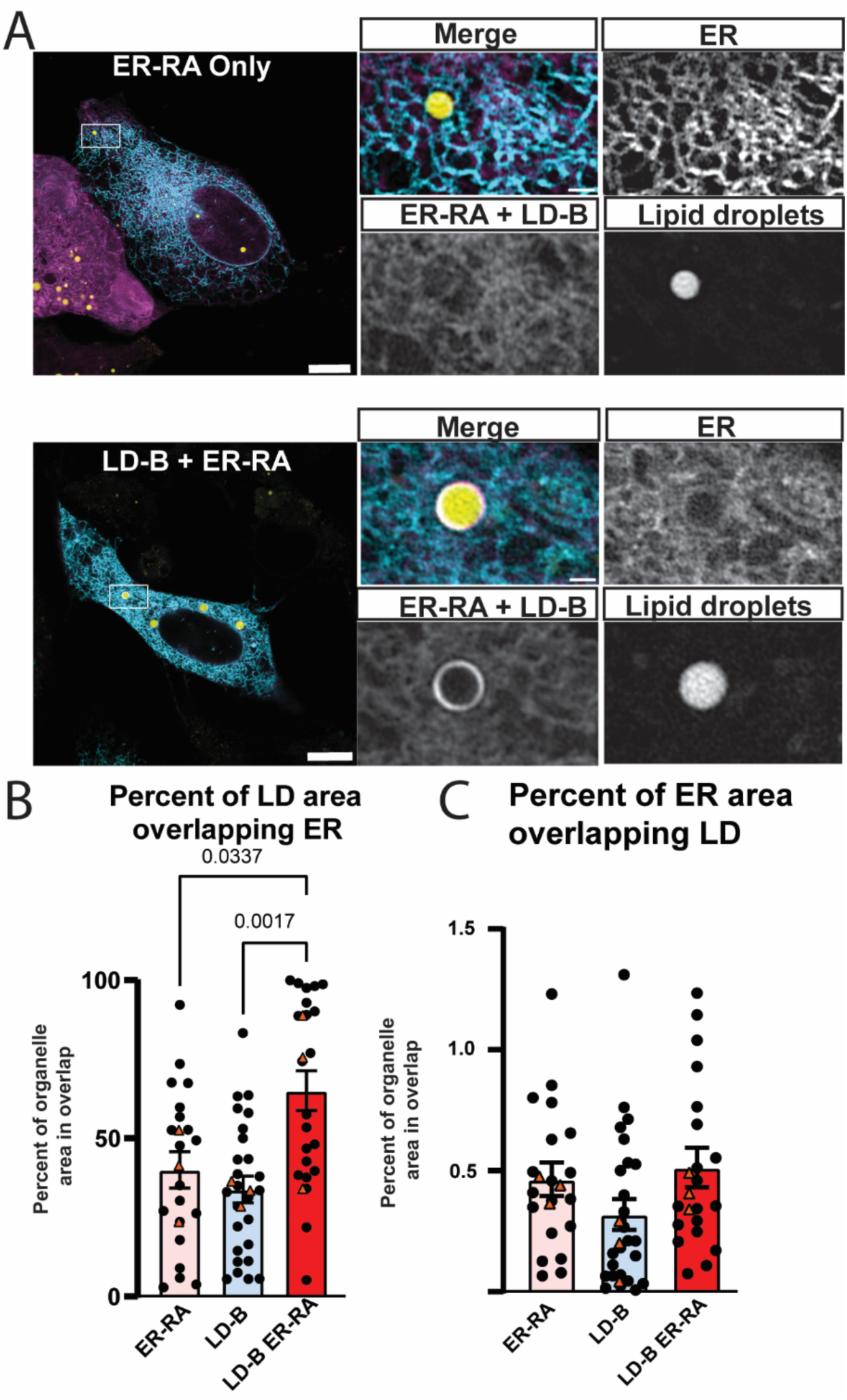
Contact-FP can induce ER-LD contacts. A) Micrographs showing transfection of ER-RA with LD-B increases overlap of ER and LD in U-2 OS compared to ER-RA transfected alone. BFP-KDEL used to label ER, BODIPY493 used to label LDs. B) Quantification of overlap of ER-LD objects, showing percent of each category of objects’ area touching the opposing objects’ segmented area. n>25 cells per condition across 3 replicates. Each point represents a single cell. Medians from each replicate are shown with orange triangles. Error bars represent standard error of the mean (SEM). Scale bars: 10 µm.

We then tested whether artificial tethering of LDs to the ER using ddFPs is sufficient to affect the morphology of LDs (Figure 3A). Strikingly, we found that inducing ER-LD contacts with ddFPs increased LD size compared to untransfected cells or cells transfected with the ER-RA or LD-B monomers individually (Figure 3B) and led to a modest but not statistically significant decrease in LD count compared to controls (Figure 3C). This increase in LD size did not lead to an increase in total LD amount, measured as LD percentage of cell area (Figure 3D), suggesting that induction of MCSs via ddFPs modifies the distribution of cellular lipid in LDs. Interestingly, the effect of artificially tethering LDs to the ER on LD morphology differs from inducing increased LD biogenesis through supplementation with the monounsaturated dietary fatty acid oleic acid (OA). OA supplementation is a well-established tool to increase LD biogenesis.

**Figure 3:**
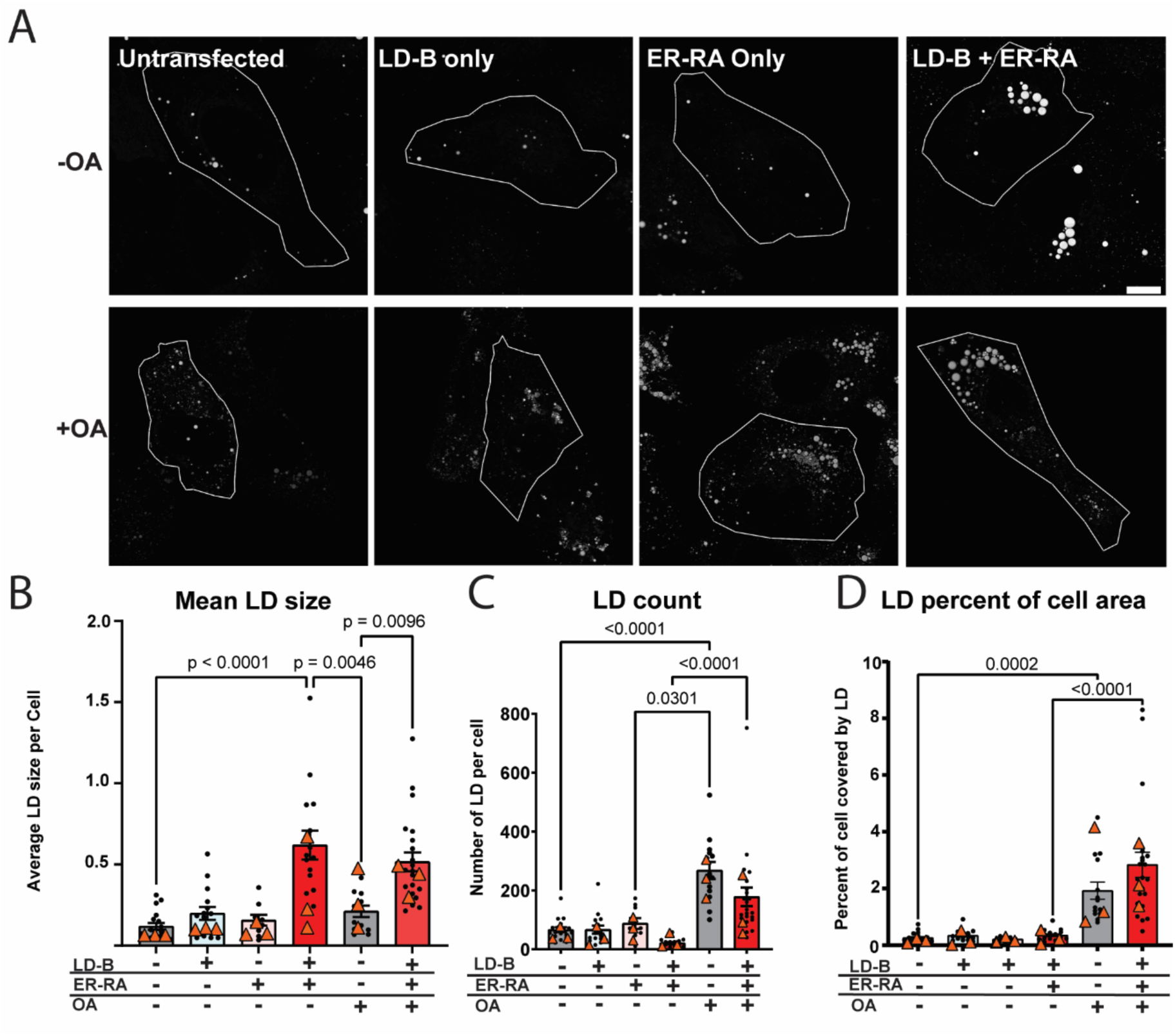
Artificially inducing ER-LD contact sites increases LD size. A) Micrographs displaying BODIPY493 channel of untransfected cells, cells transfected for two days with each Contact-FP monomer alone or with the combined Contact-FP monomers with/without oleic acid treatment as specified in the graph. Solid line designates cell area. B) Quantification of LD size, percent area of cell covered by segmented LD objects and number of LD as quantified in FIJI. n>26 cells per condition across 3 replicates. Each point represents a single cell. Medians from each replicate are shown with orange triangles. Error bars represent SEM. Scale bars= 10 µm

Regardless of OA supplementation, however, inducing ER-LD contacts with ddFPs increased LD size compared to untransfected controls (Figure 3B). ddFP-transfected cells were still responsive to OA, increasing LD number and percent of cell area (Figure 3C-D), but not size (Figure 3B), suggesting that inducing ER-LD contact sites modifies LDs by a different mechanism than OA supplementation. Taken together, we found that inducing ER-LD MCSs not only increased ER-LD colocalization but also increased LD size without modifying LD percent of cell area. This suggests that manipulation of ER-LD MCSs modifies the distribution of cellular lipid in LDs without increasing the total amount of cellular lipid stored in neutral lipids or increasing the rate of new LD biogenesis. ER-LD MCS proteins have long been known to modify the number and morphology of LDs. In the absence of ER-LD contact site proteins such as seipin, LDs can spontaneously form, but the control of their size and number is affected, leading to a population composed of supersize and very small LDs (Salo, et al, 2016; Wang et al, 2016; Salo et al, 2019). Our data suggest that artificially inducing ER-LD MCSs may be sufficient to modify the way lipid is distributed into existing LDs, shifting the average amount of lipid incorporated into existing LDs. While ddFP-transfected cells treated with OA have larger LDs than untransfected cells, they are still responsive to OA treatment in terms of LD count and percent area (Figure 3B-D). These data together suggest that inducing ER-LD MCSs with ddFPs shifts LD size, causing the cell to store more neutral lipid per LD without perturbing the ability of the cell to respond to fatty acid stimulation and undergo normal LD biogenesis.

### Exogenously increasing ER-LD contacts leads to increased proximity, but not full recruitment, of TAG synthesis enzyme DGAT2 to LDs

We hypothesized that our constructs increased average LD size either through recruitment of lipid synthesis enzymes from the ER and/or movement of lipids into existing ddFP-positive LDs. We next sought to discern whether our artificial membrane tethers could perform either of two essential functions of ER-LD contact site proteins: facilitating protein recruitment from the ER or allowing for the movement of ER-localized neutral lipid into existing LDs. One possible mechanism by which our ddFP system might increase LD size is by increasing local synthesis of neutral lipids at ddFP-positive LDs via recruitment of lipid synthesis enzymes. 90% of TAG synthesis in murine models has been shown to be dependent on one of two rate-limiting enzymes, DGAT1 or DGAT2. DGAT2 is known to diffuse onto the LD surface upon stimulation of LD biogenesis and, like our ddFP system, overexpression of DGAT2 leads to increased size of a subpopulation of LDs (Wilfling et al, 2013; Kuerschner et al, 2007; Stone et al, 2009, Stone et al, 2004). Given the similarity in phenotype and the known diffusion of DGAT2 onto LDs, our ddFP system might increase the local concentration of DGAT2 on ddFP-positive LDs, increasing local TAG synthesis at these ER-LD MCSs and increasing overall LD size.

Therefore, to test whether artificially induced ER-LD contact sites can recruit proteins from the ER, we examined the recruitment of DGAT2 to LDs in ddFP-transfected cells. Importantly, under conditions of supplementation with OA, DGAT2 can associate with LDs and promote local TAG synthesis at the surface of the LD (McFie et al, 2011; Stone et al, 2009; Kuerschner et al, 2007). Because the weak fluorescence of ddFP dimers is not well preserved with fixation, we exogenously expressed GFP-tagged DGAT2 together with the ddFP constructs. We then compared DGAT2 localization in ddFP-transfected cells to DGAT2-GFP-only transfected controls (Figure 4A). We found an increase in normalized intensity of DGAT2-GFP around LDs in ddFP transfected cells compared to control (Figure 4B). However, our images showed a tubular ER-like morphology of DGAT2-GFP in ddFP-transfected cells, similar to that of the DGAT2-GFP only transfected cells without OA, rather than a punctate or ring-like morphology that would suggest a primarily LD localization (Figure 4A). When we supplemented DGAT2-GFP transfected cells with OA, we observed a more dramatic change in localization of DGAT2-GFP from an ER morphology to rings around the LD (Figure 4A). Indeed, when we treated ddFP-transfected cells with OA, we observed a very similar change in the localization of DGAT2-GFP, suggesting a more complete recruitment of DGAT2-GFP to LDs than ddFP transfection alone (Figure 4B). To distinguish between the effect of ddFP transfection and OA treatment, we compared the mean DGAT2-GFP fluorescence intensity at ddFP-positive versus -negative LDs in the same cells. Regardless of OA supplementation, mean DGAT2-GFP recruitment was higher on ddFP-positive LDs compared to their ddFP-negative counterparts within the same cell (Figure 4C).

**Figure 4:**
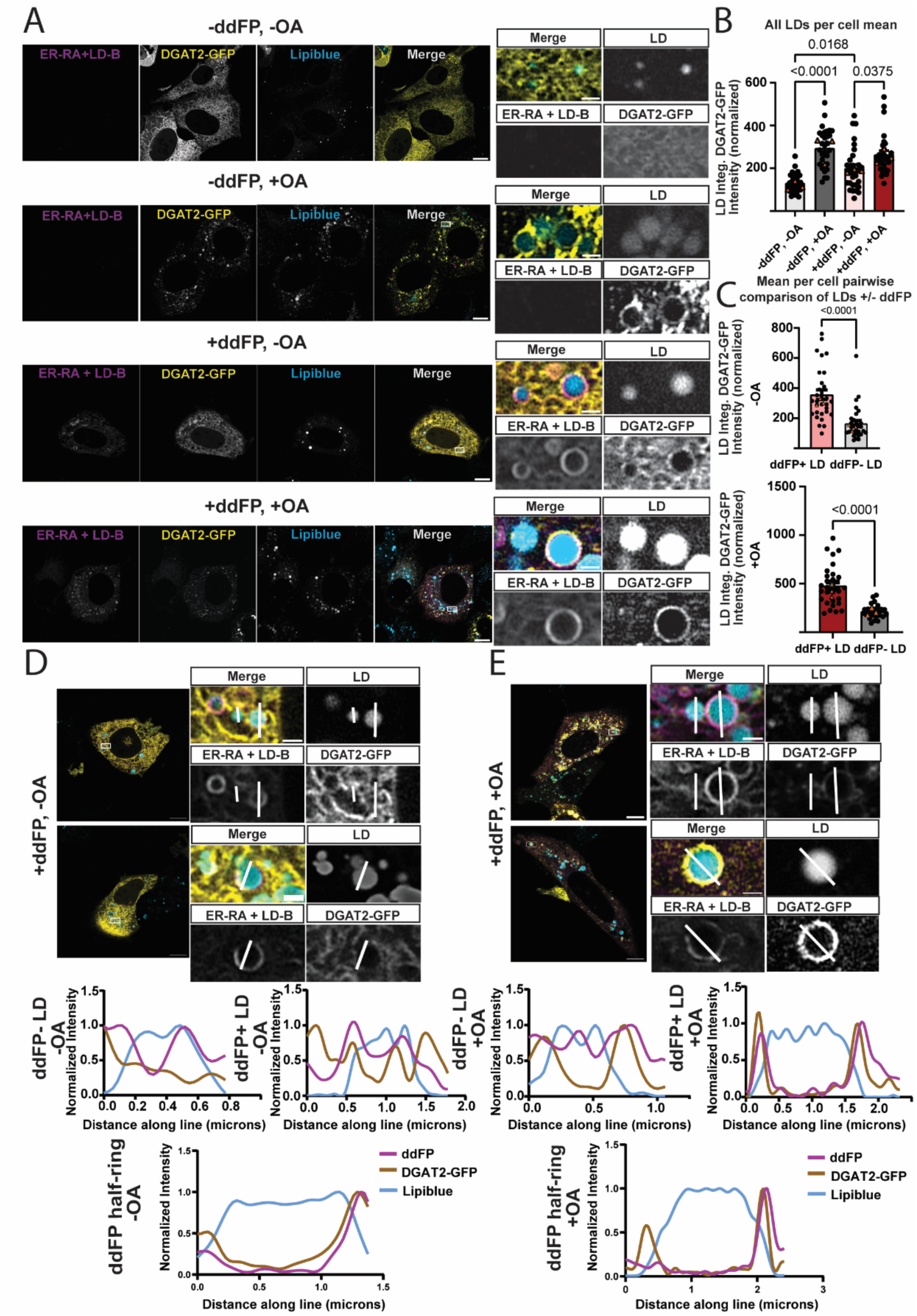
Contact-FP recruits DGAT2 to the ER-LD interface but not the LD surface. A) Micrographs showing localization of ER-RA with or without ddFP monomers and with or without 5-hour 400 uM oleic acid supplementation. DGAT2-GFP translocates from the ER to the LD surface under OA stimulation, but to the ER-LD interface with Contact-FP transfection. LDs labeled with Lipiblue dye. B) Quantification of mean integrated DGAT2-GFP intensity around LD normalized to mean DGAT2-GFP intensity within the cell under specified conditions. C) Quantification of average DGAT2-GFP integrated intensity around ddFP positive or negative LD per cell normalized to average DGAT2-GFP intensity in the whole cell with or without oleic acid. Both conditions show a significant increase in DGAT2-GFP intensity normalized with OA supplementation. D) Paired per-cell average of DGAT2-GFP normalized integrated intensity shows that within individual cells, ddFP+ LD have more DGAT2-GFP intensity relative to ddFP-, regardless of OA supplementation. n>29 cells per condition across 3 replicates. Each point represents a single cell. Medians from each replicate are shown with orange triangles. Error bars represent SEM. Scale bars: 10 µm. E-F) Line scans showing the recruitment of DGAT2-GFP to ddFP-positive, ddFP-half-ring, or ddFP-negative LDs with and without 5-hour oleic acid supplementation.

When examining ddFP-positive versus ddFP-negative LDs in DGAT2-GFP and ddFP transfected cells by linescan, relative DGAT2-GFP intensity around LDs is modestly increased at ddFP-positive LDs, but is not greater than other points further from the LD along the line (Figure 4D). When a LD is partially contacting the ER but not fully enveloped (see Figure 4D, ddFP half-ring), we observe greater local enrichment of DGAT2-GFP in the ddFP-positive areas of the LD compared to the areas of less ddFP intensity. These data suggest that the brighter fluorescence of DGAT2-GFP at the LD is higher than background but may be of similar intensities to other regions of the cells, such as ER areas not in close proximity to LDs. Thus, the quantified increase in DGAT2-GFP around ddFP-positive LDs may result from the increased proximity of ER-localized DGAT2-GFP to LDs where ER-LD MCSs are induced rather than movement of DGAT2 onto the LD surface. When we examined ddFP and DGAT2-GFP-transfected cells supplemented with OA, we saw local recruitment of DGA2-GFP to both ddFP-positive and ddFP-negative LDs (Figure 4E). However, in ddFP-positive LDs the peak DGAT2-GFP relative intensity is sharper and drops off more dramatically further from the LD, suggesting increased DGAT2-GFP recruitment to the ER-LD interface (Figure 4E).

When partial ddFP rings were observed, increased DGAT2-GFP intensity was visible at both sides of the LD only in OA-treated cells (Figure 4E). In untreated cells, DGAT2-GFP peaks were observed only on the ddFP-positive side of the LD (Figure 4D), consistent with DGAT2 being in the ER at an MCS but not on the LD surface. These results suggest that ddFP-induced contacts bring DGAT2 into closer proximity to LDs due to the increased tethering of the ER to LDs, but fail to support the full trafficking of DGAT2 from the ER to LDs.

Known ER-LD proteins such as seipin have roles in both recruiting cytosolic-facing ER proteins to the LD surface or restricting these proteins access to the LD (Salo et al, 2016). Our data indicate that while exogenous manipulation of MCSs does increase some local proximity of the key TAG synthesis enzyme DGAT2 to ER-LD MCSs, it does not facilitate a complete re-localization of DGAT2 from the ER to the LD surface in the same manner as OA supplementation, even locally at ddFP-positive LDs. This suggests that while DGAT2-GFP may be enriched at ER-LD MCSs at ddFP-positive LDs, these artificial contact sites are unable to fully traffic ER proteins to the LD surface. Indeed, previous studies showed that key trafficking machinery is required for ERTOLD targeting of DGAT2 and other cargoes to LD (Song et al, 2022; Malis et al, 2024). These data together suggest that, on their own, artificial tethers are sufficient to increase the local concentration of DGAT2 in the ER close to LDs, which may allow for some of their products to more easily incorporate into existing LDs, but the tethers are not able to facilitate the change in localization of ERTOLD cargoes to LDs.

### Artificial ER-LD tethering with ddFPs funnels fatty acids into existing LDs

Because artificially tethering LDs to the ER was not sufficient to recruit DGAT2 from the ER to the LD surface for local TAG synthesis to drive LD expansion, we hypothesized that inducing ER-LD contacts may facilitate the growth of LDs by increasing the movement of lipid from the ER and nascent lipid droplets connected to the ER into existing ddFP-positive LDs. We also hypothesized that our artificial tether could increase the rate of fatty acids moving directly from the ER/nascent LDs into existing LDs. This could counteract the lipid partitioning functions of other ER-LD proteins and facilitate ripening by biasing fatty acid redistribution into ddFP-positive LDs by their retained access to the ER/nascent LD pool of neutral lipid. To test this hypothesis, we expressed ER-LD ddFPs and labelled existing LDs overnight with Lipiblue. Then, an hour prior to imaging, we treated cells with the fluorescent fatty acid analogue FL-C12 and quantified its incorporation into LDs (Figure 5A). Strikingly, we found that ddFP-positive LDs were larger and showed increased incorporation of FL-C12 compared to ddFP-negative LDs within the same cell (Figure 5B-D), suggesting that ddFP-induced contacts increase the movement of cellular lipid into existing LDs. Taken together, our data suggest that artificial tethering of LDs to the ER is sufficient to induce the movement of cellular FAs, but not lipid synthesis enzymes such as DGAT2, to existing LDs (see model in Figure 5E). These data suggest that direct movement of fatty acid from the ER into existing LDs in contact with the ER can be facilitated by our tethers and that this movement of fatty acid is likely the driving force behind the increased size phenotype. ddFPs likely increase the incorporation of FAs from the ER or nascent LDs in proximity to the ddFP positive LDs. Proteins on the LD surface are known to decrease surface tension and increase their stability (Thiam et al, 2013). Additionally, when oil in water emulsions are in proximity, it is energetically favorable for small emulsions to fuse to reduce surface tension in the process of ripening (Thiam et al, 2016). We propose that ddFP-positive LDs increase in size and preferentially incorporate new lipid because they are both stabilized by the decoration of the monomers on the LD surface and because the anchoring of these LDs to the surface of the ER facilitates the increased ripening of these LDs by providing them access to the neutral lipid accumulation in the leaflets of the ER. However, given the observation that our system does not significantly decrease LD number or percentage area, and still allows for the increase in LD number and area with OA stimulation, artificial tethers likely cannot fully divert all exogenous lipid into existing LDs and still permit the formation of some new LDs.

**Figure 5:**
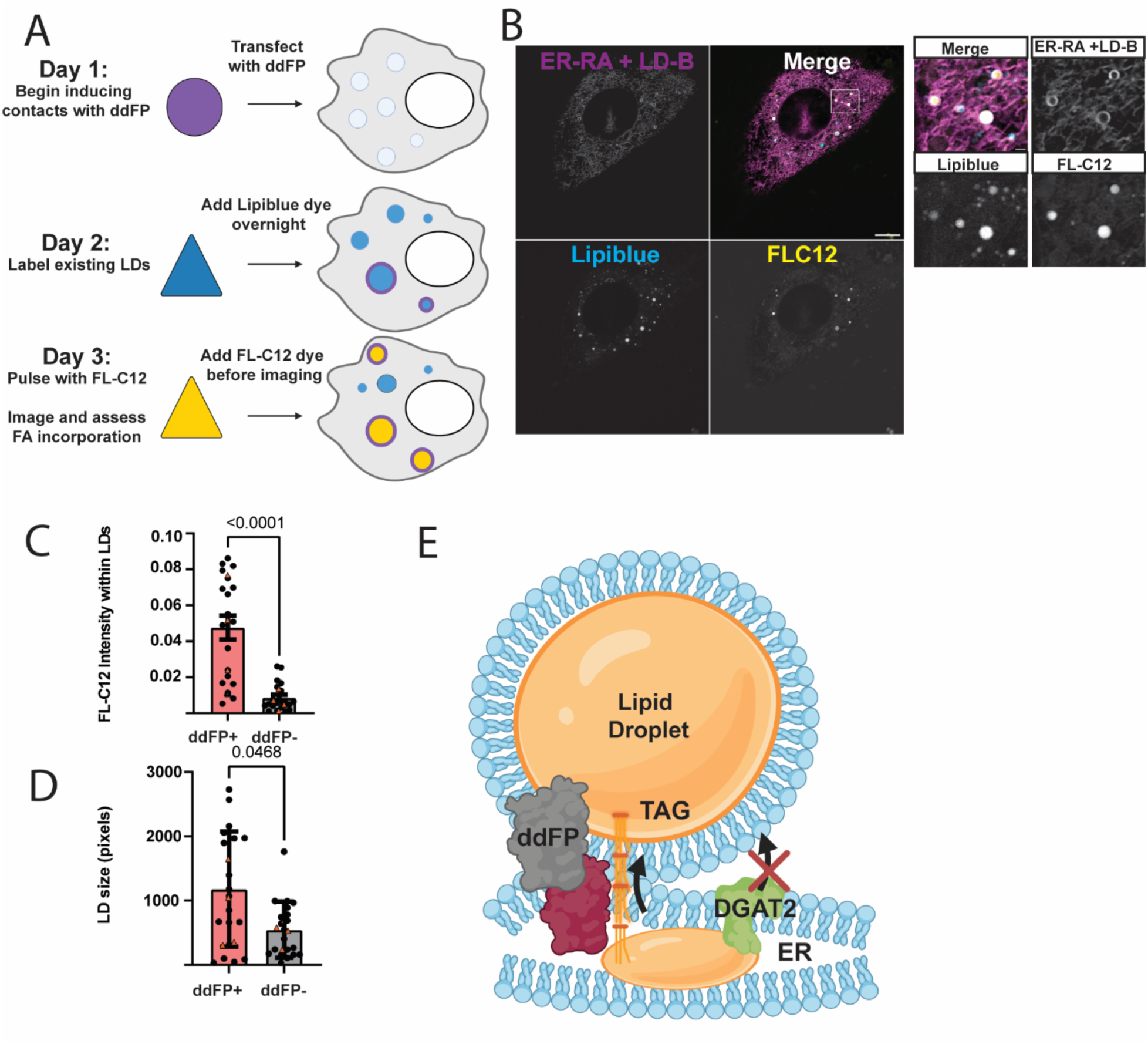
Contact-FP channels lipid into existing LDs. A) Schematic of experimental design. Cells were transfected with ER-RA and LD-B ddFPs (purple) and 24 hours before imaging, supplemented with LipiBlue dye (blue) to label LDs. FL-C12 (yellow) was added to media one hour prior to imaging to measure fatty acid uptake into existing ddFP-positive versus ddFP-negative LDs. B) Micrographs, and C-D) quantification of the experiment described in (A). ddFP-positive LDs are both larger and incorporate more FL-C12 compared to ddFP-negative LDs in the same cells. n>29 cells per condition across 4 replicates. Each point represents a single cell. Medians from each replicate are shown with orange triangles. Error bars represent SEM. Scale bar: 10 µm. E) Proposed model for the mechanism by which Contact-FP increases LD size.

## Discussion

In this study, we demonstrate the use of dimerization-dependent fluorescent proteins (ddFPs) to exogenously manipulate ER-LD MCSs. Using these tools, we interrogated which functions of ER-LD contact site proteins can be recapitulated using artificial tethers and which properties of ER-LD MCSs are dependent on specific protein structure and function. After confirming the localization of the ER-RA and LD-B ddFP monomers that we employed in this study, we found that these two constructs together efficiently and specifically label ER-LD MCSs (Figure 1). Importantly, our biosensors can distinguish LDs fully wrapped in a ring of ER, participating in small areas of contact, or harboring no ER-LD contacts at all (Figure 1D). While the current study is devoted to the exogenous manipulation of ER-LD MCSs, these data reinforce the ability of these probes to detect ER-LD contact sites (Miner et al, 2024). Given the selectivity of ddFPs in identifying LDs participating in ER-LD MCSs, we believe this sensor can continue to be useful in providing a microscopy approach to discriminate between LDs participating in MCSs, and for visualizing MCS dynamics in live cells. ddFPs can provide an efficient readout to detect differences between LDs that are involved in ER-LD MCSs versus those that are not, in response to various perturbations. While the interaction between ddFP monomers is reversible and can allow for dynamic association and dissociation between organelles at low expression levels, a caveat is that ddFPs can stabilize and induce contacts when transfected in high amounts (see Williams et al, 2025). At the concentrations transfected in this study, ddFP expression increases ER-LD MCSs, providing a use-case for this tool to manipulate the interactions between membrane-bound organelles (Figure 2). We have here presented a way to leverage ddFPs to study the effect of artificial ER-LD MCSs on LD morphology.

In this study, we sought to characterize whether ddFP artificial tethers could perform the functions of known ER-LD tether proteins. Because we observed an increased LD size phenotype (Figure 3), we sought to distinguish how artificial tethers increase lipid incorporation into LDs at MCSs, either indirectly through recruitment of TAG synthesis enzymes or directly through movement of ER-localized neutral lipid into tethered LDs.

We found a modest but incomplete recruitment of the TAG synthesis enzyme DGAT2 to ER-tethered LDs, suggesting that the artificial tether can increase proximity but not fully relocalize proteins from the ER to LDs (Figure 4). In contrast, we saw direct preferential incorporation of exogenous lipid into tethered LDs, indicating that artificial tethering can direct lipid from the ER or nascent LDs into existing LDs (Figure 5).

Overall, our study highlights the potential of manipulating MCSs to influence organelle biology and behavior. We have found a new use for ddFPs to modify the lipid distribution within LDs through manipulation of ER-LD MCSs. We found that our artificial tethering system promotes lipid transfer to LDs and promotes recruitment of proteins to ER-LD MCSs without allowing for full recruitment of these proteins onto the LD surface. This highlights an important distinction in function of ER-LD MCSs between those that facilitate increased lipid vs. protein translocation. These data also support the importance of tether stoichiometry in mediating the downstream effects on LD size, for example as seen in seipin degron systems that lead to an increase in LD size (Salo et al, 2019). Further, these data suggest a persistent function of ER-LD MCSs in LD growth after LD biogenesis. While ER-LD MCSs that occur after LD biogenesis have been suggested to mediate protein trafficking, our exogenous system also suggests a function for these contacts in increasing and maintaining LD size over time. Given that some mammalian LDs retain contact with the ER while others do not, the persistence or lack of ER-LD MCSs may suggest a different function of LD sub-populations as longer or shorter-term structures. Our data show that longer term contact with the ER allows increased lipid channeling to the LD over time, suggesting a distinct role for this sub-population that may have implications for energy storage and cellular responses to lipid stressors. While our study focused on visualizing and inducing contacts in U-2 OS cells, we envision using ddFPs to manipulate LDs in other systems in the future. In cell types such as adipocytes and hepatocytes, visualizing and investigating how manipulating ER-LD contacts affects cellular function, differentiation and disease progression may provide valuable insight.

## Materials and Methods

### Plasmids

Contact-FP plasmids were generated by the Sarah Cohen lab and are available on Addgene (Miner et al, 2024). Used in this study were Cyto-RA (Addgene #209860), Cyto-B (Addgene #209858), ER-RA (Addgene #209872) and LD-B (Addgene #209861). TagBFP-KDEL construct was a generous gift from Gia Voeltz (University of Colorado; Friedman et al., 2011; Lee et al., 2020); DGAT2-GFP was a generous gift from Koret Hirschberg (Tel Aviv University).

### Cell Culture

U-2 OS cells were procured from UNC Tissue Culture Facility and maintained in Dulbecco’s Modified Eagle Medium (Corning, MT15017CV) with the addition of 10% fetal bovine serum (VWR # 97068-085), penicillin/streptomycin (Corning MT30002CI; working concentration 100IU/100ug/mL) and 4mM glutamine (Corning, MT25005CI). Cells were seeded onto 8-well chambered cover glass (Cellvis # C8-1.5H-N) coated with 10ug/mL fibronectin (MilliporeSigma #FC010-1MG) for imaging. Imaging media was prepared using DMEM without phenol red (Corning 21041025) with 10% FBS and 4mM glutamine and without penicillin/streptomycin.

### Transfection

Cells were transfected with indicated plasmids using Lipofectamine 3000 (Invitrogen #L3000008) for 16 hours in base media according to manufacturer instructions, using OptiMEM (Gibco #31985070) as vehicle for transfection. Unless otherwise specified, 25 ng of LD-B, 50 ng of ER-RA and 10 ng of BFP-KDEL were transfected per 8 well chamber slide chamber. Base media was changed after 16 hours and cells were imaged 48 hours after transfection. For oleic acid treatment, cells were treated with 400 uM sodium oleate (Sigma #O7501) for 5 hours or overnight. Cells were washed once with imaging media and supplemented with either 500 nM Lipi-Blue dye (Dojindo #LD01) one hour prior to imaging or 200 nM BODIPY493 (Invitrogen #D3922) 30 minutes prior to imaging in imaging media. For FL-C12 supplementation experiments, cells were labelled overnight with 500nM Lipi-Blue. 10 uM BODIPY FL-C12 (Invitrogen #D3822) was added to imaging media one hour prior to imaging.

### Imaging

Images were captured using an inverted Zeiss 800/Airyscan laser scanning confocal microscope equipped with 405, 488, 561, and 647 nm diode lasers, and Galium Arsenide Phosphide (GaAsP) and Airyscan detectors. Airyscan images were obtained using a 63×/1.4 numerical aperture objective lens, at 37 °C and 5% CO2 (Carl Zeiss, Oberkochen, Germany). Airyscan images were processed using Zen software (Carl Zeiss) with a processing strength of 6.0.

### Quantification and Statistical Analysis

Images were analyzed using CellProfiler to segment organelles and regions of interest (ROIs), to measure fluorescent intensities within ROIs and to identify adjacent objects, including LDs, ER, ddFPs and DGAT2-GFP(Stirling et al., 2021). Total cell fluorescence intensities were calculated from image channels within manually identified cell objects with nuclei excluded. Colocalization measurements of ER and LD fluorescence channels were calculated by segmenting each object and measuring percent of object touching the opposing organelle’s segmented object within the manually identified cell area. Lipid droplets and ddFP objects were thresholded and segmented within the resulting objects. Lipid droplet masks were expanded by 3 pixels to allow for capture of the area surrounding the object to identify adjacent ddFP or DGAT2-GFP signal. ddFP-positive lipid droplets were identified through running the IdentifyObjectNeighbors module on expanded lipid droplet objects and the segmented ddFP objects and lipid droplet objects with 1 or more ddFP object neighbors and a percent touching of adjacent ddFP objects of >0 were designated as ddFP positive. Intensity around lipid droplets was used by masking the expanded lipid droplet object with the original lipid droplet object and measuring intensities of the raw image within the area of these objects. For FL-C12 experiments, FL-C12 and LipiBlue channels were individually segmented and expanded by 3 pixels and fluorescent intensities were calculated within these segmented objects. Line scan data were generated in FIJI using the Plot Profile tool along the indicated ROI using the raw image intensity values and plotted in GraphPad Prism, normalizing to the highest intensity value per fluorescence channel along the line.

Lipid droplet number, size and percent area measurements under ddFP contact induction were measured using a FIJI script as described in Windham et al, 2024. In brief, individual cells were manually segmented and the area outside of each cell was excluded from segmentation with the “Clear Outside” function. To segment LDs, the BODIPY493 or Lipiblue channel was first passed through a Gaussian filter followed by a Laplacian of Gaussian (LoG) filter optimized in radius based on image intensity. Cells were autothresholded using the Otsu algorithm and “Analyze Particles” was used to measure the area and count of segmented LDs, as well as the sum total area of all LDs within the cell segmentation.

Statistics and plotting of data were performed in GraphPad Prism. Outliers were identified using ROUT with Q= 5%. Statistical analyses among groups was performed using a Mann-Whitney unpaired test for direct two group of cell comparisons, Wilcoxon test for paired comparisons of ddFP-positive versus ddFP-negative lipid droplets within the same cell, or a Kruskal-Wallis one-way ANOVA with Dunn’s correction for multiple comparisons or comparison of multiple conditions simultaneously.

## Author Contributions

V.H. Williams, G.E. Miner, and S. Cohen designed the project and experiments. V.H. Williams performed experiments and data analysis. V.H. Williams and S. Cohen wrote the manuscript. S. Cohen acquired funding and supervised the project.

## Acknowledgements

We would like to thank Gia Voeltz and Koret Hirschberg for the specified plasmids used in this study. We would also like to thank Yehonathan Malis for his feedback during the course of this study. Research reported in this publication was supported by the National Institutes of Health under award number R35GM133460. Illustrations in this study were generated using BioRender.

